# SuperTranscript: a data driven reference for analysis and visualisation of transcriptomes

**DOI:** 10.1101/077750

**Authors:** Nadia M Davidson, Anthony DK Hawkins, Alicia Oshlack

## Abstract

Numerous methods have been developed to analyse RNA sequencing data, but most rely on the availability of a reference genome, making them unsuitable for non-model organisms. De novo transcriptome assembly can build a reference transcriptome from the non-model sequencing data, but falls short of allowing most tools to be applied. Here we present superTranscripts, a simple but powerful solution to bridge that gap. SuperTranscripts are a substitute for a reference genome, consisting of all the unique exonic sequence, in transcriptional order, such that each gene is represented by a single sequence. We demonstrate how superTranscripts allow visualization, variant detection and differential isoform detection in non-model organisms, using widely applied methods that are designed to work with reference genomes. SuperTranscripts can also be applied to model organisms to enhance visualization and discover novel expressed sequence. We describe Lace, software to construct superTranscripts from any set of transcripts including de novo assembled transcriptomes. In addition we used Lace to combine reference and assembled transcriptomes for chicken and recovered the sequence of hundreds of gaps in the reference genome.

## Background

High throughput sequencing has revolutionized transcriptomics because it allows cDNA sequence to be read and expression levels quantified using a single, affordable assay[1, 2]. RNA sequencing (RNA-seq) can examine expression at the gene level as well as infer transcript abundances and differential isoform usage. Alternative splicing can alter gene function and contributes to the overall transcriptional diversity in eukaryotes[3, 4]. In addition, RNA sequencing has the power to detect variation in expressed sequence, such as single nucleotide variants[5], post-transcriptional editing[6] and fusion genes[7]. Our knowledge of the transcriptome of model organisms has matured through projects such as ENCODE[4] and we now have robust and well established methods for RNA-Seq data analysis[8]. Most of these methods use the accurate reference genomes and annotations now available for model organisms.

For non-model organisms, however, reference genomes are generally not available. Instead, an experiment specific transcriptome can be built from RNA-seq data through de novo transcriptome assembly[9]; a process designed to reconstruct the full length sequence of each expressed transcript. However down-stream analysis of the data remains challenging. With the exception of a few recent methods, such as RNA quantification with Kallisto[10], Salmon[11] and RSEM[12], most analytical approaches for RNA-Seq are designed to work with a reference genome rather than transcriptome. Those methods compatible with a reference transcriptome often rely on accurate gene-models, which are not necessarily produced through de novo assembly of short read data. In addition, visualisation of read coverage across a gene, which is common for the exploration, curation and communication of the analysis results, is impossible using a reference transcriptome. At best, reads may be mapped and visualised against a representative transcript from each gene, such as the longest isoform, but a significant proportion of the gene sequence can be missed (Supplementary Table 1, Supplementary Figure 1).

Here we propose an alternative representation for each gene, which we refer to as a superTranscript. SuperTranscripts contain the sequence of all exons of a gene without redundancy (Figure 1A). They can be constructed from any set of transcripts including de novo assemblies and we have developed a python program to build them called Lace (available from https://github.com/Oshlack/Lace/wiki). Lace works by building a splice graph[13] for each gene, then topologically sorting the graph using Kahn’s algorithm[14] (Figure 1B). Building superTranscripts is a simple post-assembly step that promises to unlock numerous analytical approaches for non-model organisms (Figure 1C).

**Figure 1.**
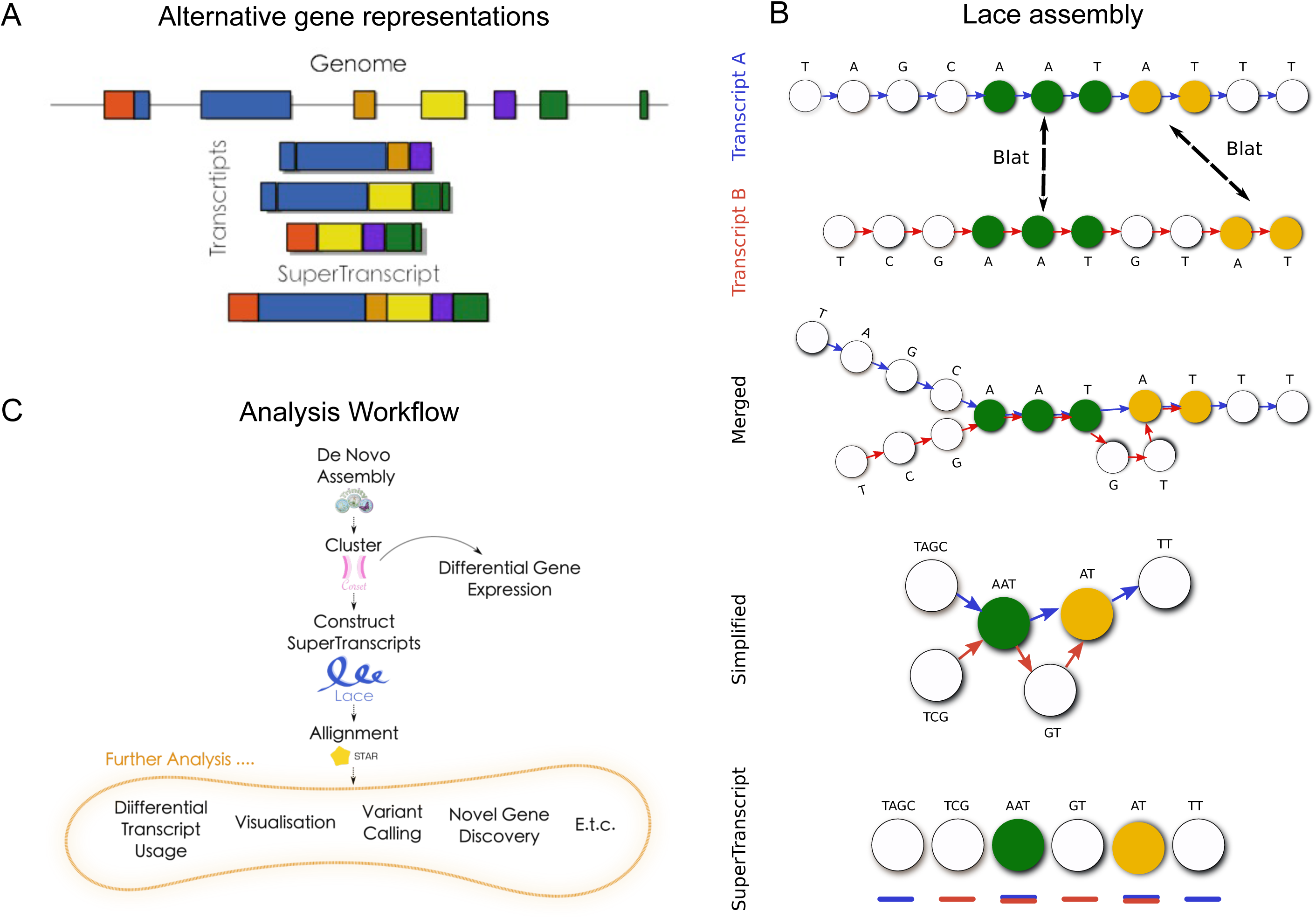
A) A gene in the genome (top) and its corresponding transcripts (middle) compared to the superTranscript for the same gene (bottom). Colours indicate superTranscript blocks. B) A schematic diagram showing the steps in Lace’s algorithm. A superTranscript (at the bottom) is built from transcripts A and B (blue and red respectively). For each transcript, Lace builds a directed graph with a node for each base. Transcripts are aligned against one another using blat and the nodes of shared bases are merged. Lace then simplifies the graph by compacting unforked edges. The graph is topologically sorted and the resulting superTranscript annotated with transcripts and blocks. C) The general workflow we propose for RNA-Seq analysis in non-model organisms. Reads are de novo assembled, transcripts clustered into genes, superTranscripts assembled using Lace and reads aligned back. Here we use Trinity, Corset and STAR as our assembler, clustering program and aligner respectively, but equivalent tools could also be used.

Although superTranscripts do not necessarily represent any true biological molecule, they provide a practical replacement for a reference genome. For example, reads can be aligned to the superTranscriptome using a splice aware aligner, and subsequently visualized using standard tools such as IGV[15]. Quantification can also be performed with existing software by counting the reads that overlap superTranscript features. In non-model organisms we further demonstrate using superTranscripts to call variants and we show that we can accurately detect differential isoform usage. We also demonstrate applications of superTranscripts to model organisms. Specifically, we combined a reference and de novo assembled transcriptome into a compact superTranscriptome using chicken RNA-seq data, to allow identification of novel transcribed sequence. We found conserved coding sequence in over 500 genes that was missed in the current chicken reference genome, galGal5.

## Results and discussion

### Lace constructs superTranscripts

SuperTranscripts can be built from any set of transcripts, including de novo assembled transcripts, using an overlap assembly method. We have implemented this algorithm in an open source Python program called Lace (available from https://github.com/Oshlack/Lace/wiki). The Lace algorithm takes two input files: (1) a set of transcript sequences in fasta format, and (2) a text file with the clustering information that groups each transcript into a gene or cluster. Lace outputs a fasta file of superTranscript sequences and a gff file with their annotation. The Lace assembly is conceptually described in Figure 1B and includes the following steps:

1. For each gene, all pairwise alignments between transcripts are performed using BLAT[16].
2. The BLAT output is parsed to determine the sequences common to each transcript pair.
3. A single directed graph is constructed per gene, where each node is a base in one of the transcripts and the directed edge retains the ordering of the bases in each transcript. Bases from overlapping sequence are merged based on the BLAT output.
4. The graph is simplified by concatenating sequences of nodes along non-diverging paths. Any cycles in the graph are detected, for each cycle the node of the cycle which contains the fewest bases is selected and duplicated. The outgoing edges from the selected node are re-routed to the duplicate node and the cycle broken (Supplementary Figure 2). This method was inspired by that used in Pezner et al.[17]. This creates a Directed Acyclic Graph.
5. The nodes are topologically sorted (each node becomes a string of bases from the original graph) using Khan's algorithm, which gives a non-unique sorting of the nodes.
6. Finally, Lace will annotate each superTranscript with blocks either by using the graph structure itself or alternatively using a script called Mobius (packaged with Lace) which infers the annotation from spliced junctions discovered when mapping reads to the superTranscript.

One of the advantages of Lace is that it can produce superTranscripts from any combination of transcripts and is compatible with any transcriptome assembler. However Lace relies on information for clustering transcripts into genes. This can be achieved with our previously published method, Corset[18] (Figure 1C).

Lace’s running time is primarily limited by the speed of the BLAT alignments, however, for genes with a large number of transcripts, processing the splicing graph is significant slower. For this reason, Lace uses only the first 50 transcripts from each gene by default. In practice, this only affects a small number of genes for most assemblies. Typically, constructing superTranscripts for an entire de novo assembly on eight cores takes approximately 0-8 hours on a linux cluster and uses up to 4 Gb of RAM, depending on the size of the input transcriptome.

### Application of Lace and superTranscripts to non-model organisms

SuperTranscripts allow a broad range of RNA-Seq analyses to be performed on non-model organisms using standard software that has been designed to work with reference genomes. To demonstrate, we analysed human RNA-Seq data without the use of the reference genome or transcriptome. First we assembled transcripts with Trinity[20] then clustered transcripts into genes using Corset[18] and subsequently built superTranscripts with Lace. Using these superTranscripts as a reference we then aligned reads back to the superTranscripts using STAR[21]. This approach allowed us to perform a variety of analysis and visualisation (Figure 1C).

### Detecting variants in non-model organisms

In model organisms, variant calling can be performed on RNA-seq data using the established GATK Best Practices workflow for RNA-Seq. Here we demonstrate that using superTranscripts as a reference allows variant calling to be performed in non-model organisms using the same pipeline with similar performance. In addition, called variants can be easily inspected in IGV for the first time (Supplementary Figure 3).

In order to demonstrate variant calling from RNA-seq data using the assembled superTranscripts as the reference we utilised RNA-Seq from Genome in a Bottle (GM12878). We called variants using the GATK RNA-seq variant calling pipeline and compared them to known variants reported by the Genome in a Bottle Consortium [22]. Specifically we took high quality heterozygous SNPs with a read coverage of 10 or more that were detected in our SuperTranscript analysis (25,788 SNPs, Supplementary Table 2). Reported homozygous SNPs were removed because they are likely to be false positives of the assembly or alignment. True homozygous SNPs should be assembled into the reference and are therefore not detectable. Next, we aligned each superTranscript back to the human genome with BLAT[16] to determined the SNP position in the genome. We then examined SNPs in the high confidence call region for Genome in a Bottle (16,483 SNPs). 82% (13,489 SNPs) were true positives reported by the Genome in a Bottle Consortium. The precision lifted to 92% if repeat regions of the genome were excluded (Supplementary Table 2).

Next we assessed how well superTranscripts compared against using the genome as a reference for calling variants. On the same dataset with the same filtering, we found that the genome-based approach gave similar results to using a superTranscript derived from de novo assembly. More true positives were reported for the genome approach (15,014 compared to 13,489), but the precision was lower (80% compared to 82%). 93% of true positives and 40% of false positives that were reported by the superTranscript approach were also reported for the genome-based approach. These results suggest that the accuracy of detecting variants in non-model organisms using superTranscripts is similar to the accuracy of detecting variants from RNA-seq in model organisms.

Finally, we validated our approach against KisSplice [23, 24], an alternative method for SNP and indel detection in non-model organisms. KisSplice performs local assembly of RNA-Seq reads and detects variants directly from the De Bruijn graph. KisSplice reported 33,252 SNPs of which 23,894 were located in the high confidence call region for Genome in a Bottle and 12,597 were true positives (Supplementary Table 2). Note that we did not filter KisSplice results for read coverage or allele balance, so cannot compare the precision against superTranscripts. However, we found the recall of KisSplice even without filtering was lower than that of superTranscripts with filtering (12,597 compared to 13,489).

### Differential isoform usage in non-model organisms

While methods exist for detecting differential gene expression in non-model organisms (such as Corset [18]), the procedure for detecting differential isoform usage is less well defined. When a reference genome is available, a common method for detecting differential isoform usage is to first align reads to the genome then count the number of reads that overlap exons for each sample (for example using featureCounts[25]) and finally perform statistical testing of the count data looking for differential exon usage using methods such as DEXSeq[26]. An alternative approach is to use estimates of transcript abundances from inference methods such as Kallisto and Salmon and subsequently perform a similar statistical testing method for differential isoform usage[27].

SuperTranscripts can be used in a similar way to the reference genome approach where reads are aligned to the superTranscripts instead of a reference genome. Each superTranscript is segmented into blocks which are used as the counting bins for statistical testing instead of exons. The blocks that annotate a superTranscript are defined as a contiguous sequence without splice junctions. Different isoforms are therefore represented by different combinations of blocks. Hence, a block may correspond to one exon, multiple exons, or part of an exon, depending on the splicing structure of the alternative transcripts within a gene. Lace provides two different types of block annotation. In one case, block positions are defined by forks or divergences in the splice graph (“Standard Blocks”), in the second case, blocks are defined dynamically using splice junctions detected in the reads that are mapped back to the superTranscript (“Dynamic Blocks”) (see Figure 2A).

**Figure 2.**
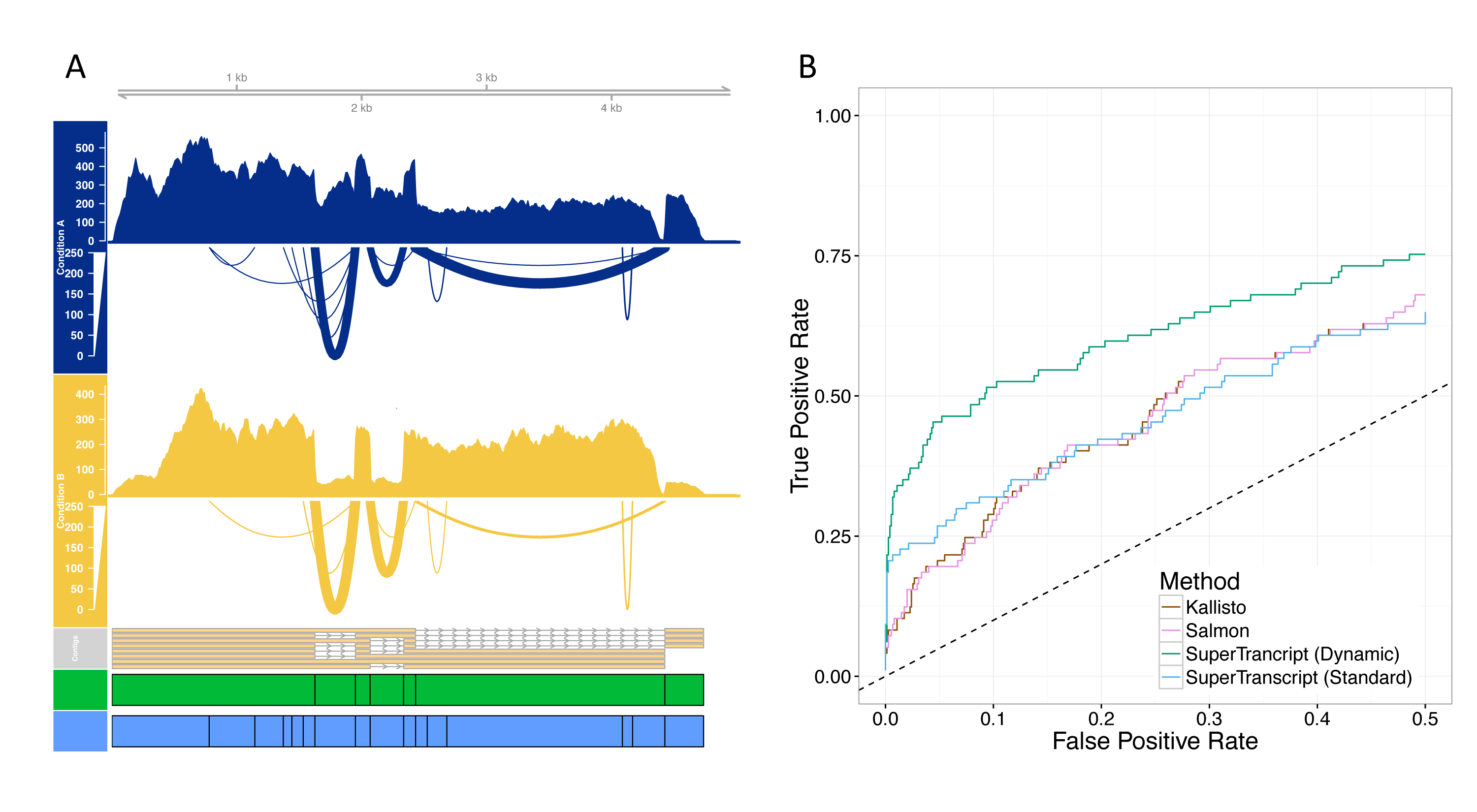
A) An example of gene visualisation for a de novo assembled transcriptome. We show the read coverage and splice junctions of Condition A (blue) and Condition B (yellow), the controls and knockdowns respectively. Differential transcript usage between two samples of different conditions can be seen in the read coverage. The two alternative block annotations for the superTranscript – Standard (green), Dynamic (blue) – are illustrated underneath. Dynamic block boundaries are located at splice junctions (defined by at least 5 spliced reads). B) ROC curve for detecting differential transcript usage using *de novo* assembled transcripts from the Trapnell et al. dataset. True and false positives are defined using a reference genome analysis (See methods).

We tested the ability of the superTranscript method to call differential isoform usage on de novo assembled data. For this we used human RNA-seq data from Trapnell *et al*.[19]. It was also analysed using a standard genome based approach for comparison. We defined genes as true positives or true negatives for differential isoform usage using the results of the genome reference based approach (see methods). We compared the accuracy of four approaches: (1) superTranscripts with standard block counts, (2) superTranscripts with dynamic block counts (3) transcript counts from Kallisto and (4) transcript counts from Salmon (see methods). We found that we had a sensitivity of 20% and specificity of 99% at FDR<0.05 using the dynamic block method. Furthermore, counting reads in superTranscript blocks performed better than transcript counts from inference methods (Figure 2B). Remarkably, dynamically defined blocks were able to detect splicing events in genes where only a single transcript was assembled resulting in an increase of sensitivity from 13% to 20% at FDR<0.05.

SuperTranscripts also provide a means of visualising differential transcript usage in non-model organisms for the first time. In Figure 2A we show differential transcript usage in ENSG00000160613 between the *HOXA1* knock-down and control groups from the Trapnell dataset. The superTranscript provides a convenient means of presenting all eight assembled transcripts, their corresponding read coverage and splicing, within a single visual. Although this type of visualisation is often taken for granted in model organisms, it is only made possible in non-model organisms by using superTranscripts as a reference.

### Combining reference and de novo assembled transcriptome

Despite the number of species with a reference genome increasing, the quality of those genomes and their annotations remain variable. Ideally, an RNA-Seq analysis could utilise prior knowledge of the gene-models available in a reference genome and annotation, whilst also extracting information about the genes from the data itself. Lace has the ability to integrate such information. It can produce superTranscripts from any source, including a combination of reference and *de novo* assembled transcriptomes. We demonstrate this idea on chicken, a model organism which we know from our previous work on chicken gonads had missing and rearranged sequence in its reference genome[28, 29].

We explored the transcriptome in chicken gonads by using Lace to assemble SuperTranscripts combining four different transcriptomes: the Ensembl annotation, RefSeq annotation, a Cufflinks[30] genome-guided assembly and a Trinity[20] *de novo* assembly (see methods). The Ensemble, RefSeq and Cufflinks transcriptomes were annotations of the galGal4 genome from November 2011. We used an older version of the reference genome so that we could validate our approach using the most recent version, galGal5 from December 2015. To construct the superTranscriptome, we first combined the genome-based annotations (Ensemble, RefSeq and Cufflinks) by merging exons and concatenating the exon sequence to build a genome-based superTranscriptome. Trinity transcripts were then aligned against the chicken genome-based and human superTranscriptome, and were subsequently assigned to gene clusters. Finally, the genome-based superTranscriptome and Trinity transcripts were assembled together with Lace. See methods for a detailed description.

The resulting superTranscriptome was compact, containing just 83Mbp (compared to almost 550 Mbp for the combined transcriptomes). However, none of the four contributing transcriptomes contained all of the sequence; 88%, 77%, 47% and 17% of bases were covered by Trinity, Cufflinks, Ensembl and Refseq, respectively. Critically, 3% (2.5Mbp) of the bases in the chicken superTranscriptome could not be found in the galGal4 reference genome. This novel sequence included superTranscripts with protein coding sequence either completely (134 superTranscripts) or partially (1332 superTranscripts) absent from the galGal4 reference genome. Figure 3A and Supplementary Figure 4 show an example of the *C22orf39* gene with a section of novel coding sequence. The novel section coincides with a known gap of approximately 100bp in the assembly of the reference genome. For most superTranscripts, sections of novel sequence typically coincided with assembly gaps (Supplementary Fig. 5A).

**Figure 3.**
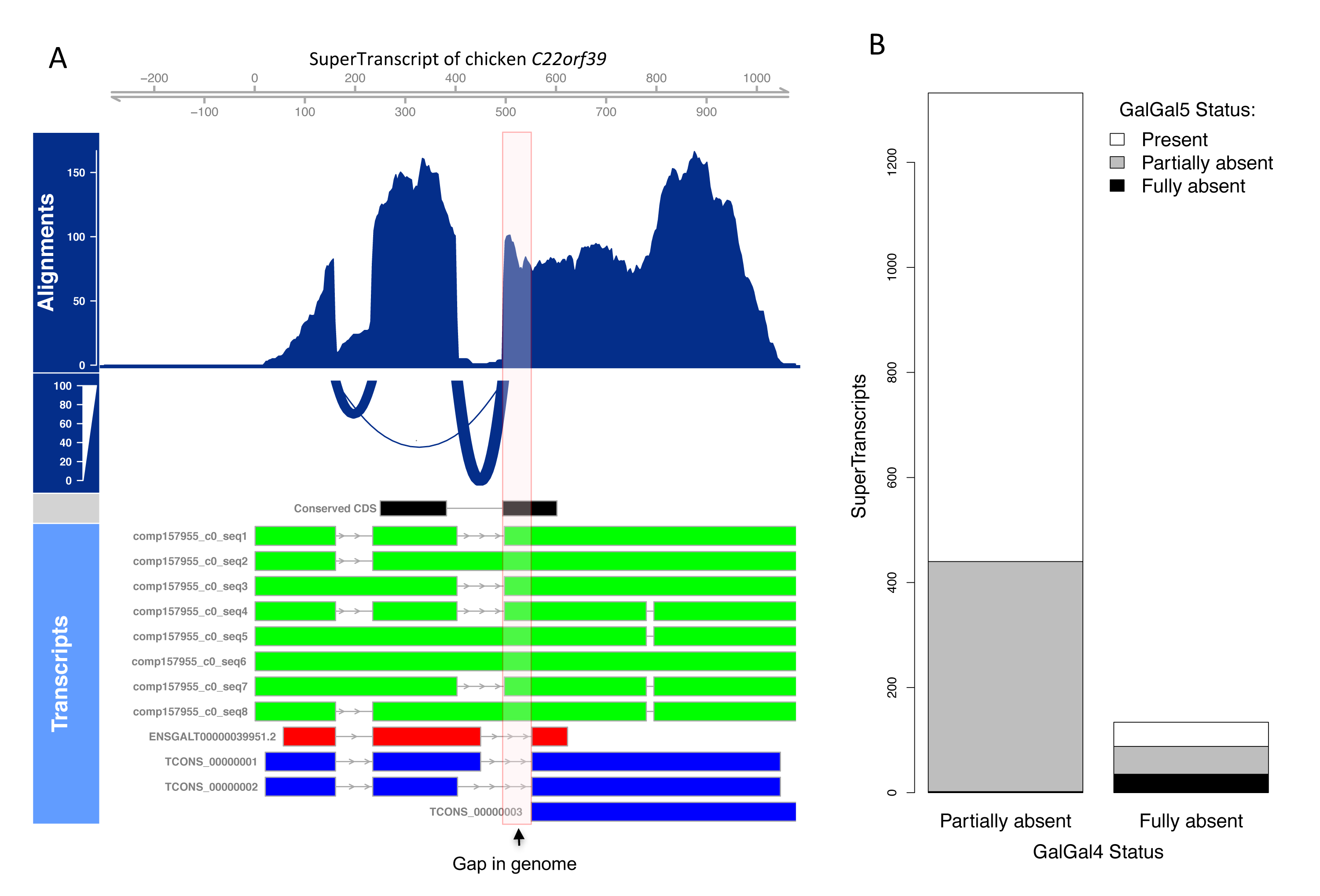
A) Reads aligned back to the superTranscript of chicken *C22orf39 (ENSGALG00000023833)*. The region shaded in red is a gap in the galGal4 reference genome. The gap is within the conserved coding sequence of the gene (black). Transcripts from Ensembl (red) and Cufflinks (blue) miss the gap sequence, whereas the Trinity assembly (green) recovers it. B) The number of superTranscripts with novel conserved coding sequence, not found in the galGal4 version of the chicken reference genome. In most cases, the superTranscript contains one or more blocks that can be found in the genome in addition to the novel blocks (partially missing), however, some superTranscripts are missing in their entirety (fully absent). Most of the novel sequence has been gained in the latest reference genome, galGal5.

To validate the novel sequence, we aligned our superTranscripts to galGal5, which contains 183 Mbp more genomic sequence than galGal4[31]. For 64% of superTranscripts with novel coding sequence in galGal4, the complete superTranscript was found in galGal5 (Figure 3B). 528 superTranscripts remain with missing sequence, including 35 that are entirely absent. This is likely because the current draft chicken genome is still incomplete[31] and approximately half the superTranscripts with novel sequence in galGal5 could be localised to regions that remain poorly assembly (Supplementary Fig 5B).

This analysis demonstrates the utility of superTranscripts and Lace to construct comprehensive transcriptome sequences in an automated way. It also highlights the major benefits of exploiting superTranscripts, even for a reasonably complete genome.

### Using SuperTranscripts in model organisms

SuperTranscripts also have a number of uses in well annotated model organisms. When a reference genome is available, we can construct superTranscripts by simply concatenating the exonic sequence of each gene rather than using Lace (we provided the superTranscriptome for human at https://github.com/Oshlack/superTranscript_paper_code). Using this superTranscriptome as a reference drastically improves visualisation because intronic sequence is excluded, giving a compact view of the mapped reads, isoform and splicing structure (Figure 4). Often this results in the ability to visualise the sequencing data (including exome data) from a whole gene simultaneously in one screen of IGV instead of having to scroll through several screens. For transcriptome data, performing read alignment with the superTranscriptome is simplified compared to a reference genome because there is less sequence and fewer splice junctions (Supplementary Table 3). In addition superTranscripts provide a convenient way of looking at the coverage and expression levels of long-read data such as PacBio or Nanopore data where each read can originate from a different isoform of the same gene (Supplementary Figure 6 shows an example for PacBio data).

**Figure 4.**
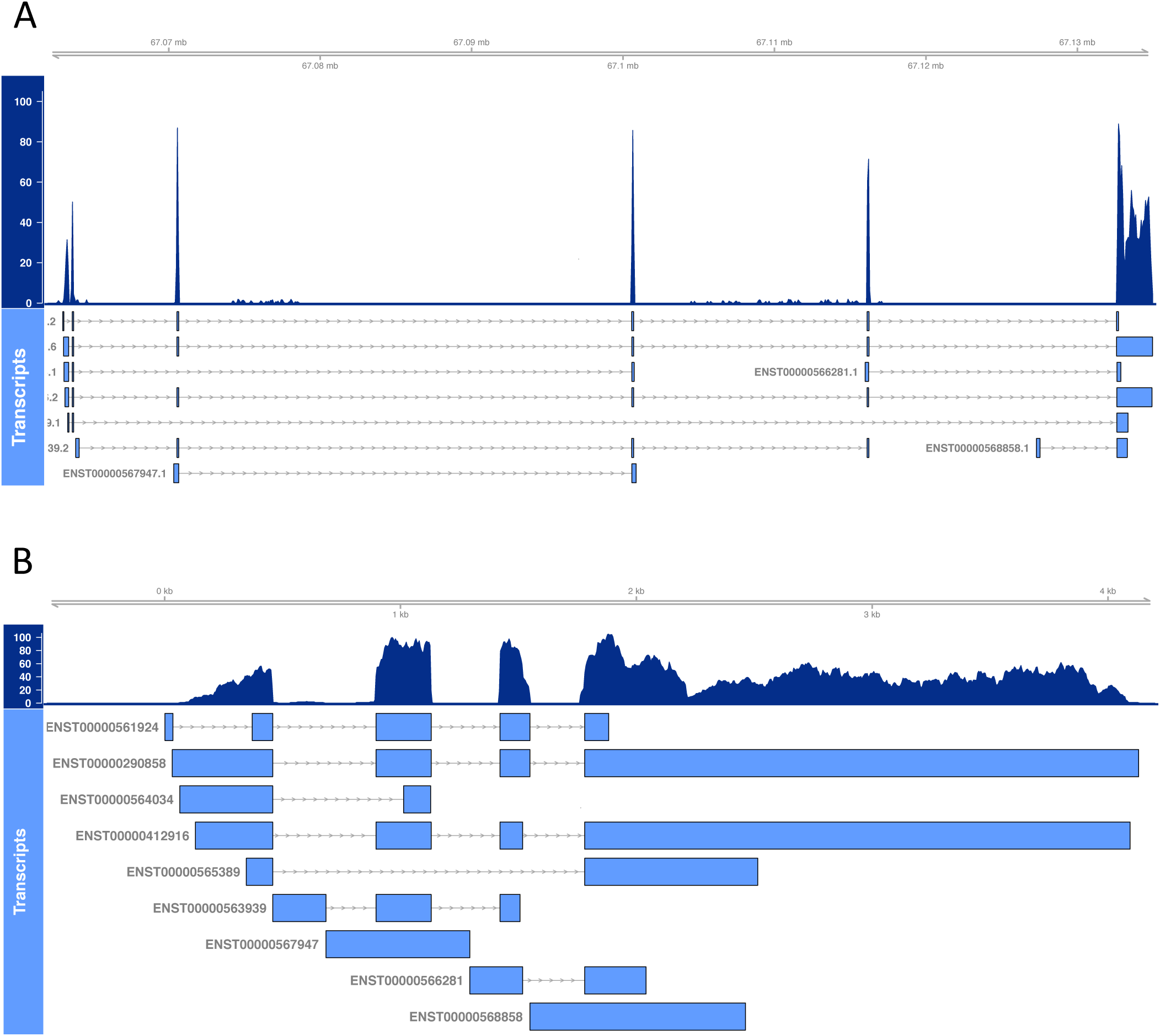
Example of the read coverage over human *CBFB* (ENSG00000067955), in A) the reference genome compared to B) the superTranscript. Transcripts are annotated below in light blue.

## Conclusions

Here we have presented the idea of superTranscripts as an alternative reference for RNA-Seq. SuperTranscripts are a set of sequences, one for each expressed gene, containing all exons without redundancy. We also introduce Lace, a software program to construct superTranscripts. Lace is unique as it is capable of assembling transcripts from any source, but existing transcriptome assemblers could also be modified to produce superTranscripts as additional output during the assembly. This would simply require the assembly graph to be topologically sorted.

Lace and superTranscripts can potentially be applied in a broad range of scenarios, some of which have been presented herein. Importantly, superTranscripts allow the visualization of transcriptome data in nonmodel organisms for the first time using standard tools such as IGV. Furthermore, superTranscripts allow differential isoform usage to be detected in non-model organisms by defining block (exon-like) structures in the transcripts and then using standard statistical testing methods such as DEX-seq. This is the first time differential isoform usage can be detected in non-model organisms using a count based approach, rather than inference methods, and we find it to be more accurate. SuperTranscripts can also be used as a reference for reliably calling variants.

A powerful, new application of superTranscripts is merging transcriptomes from a variety of sources. We demonstrate this application using a chicken transcriptome that allowed us to detect hundreds of genes containing sections of coding sequence that were not contained in the reference genome. We hypothesise that superTranscripts will have many further applications. For example, they are likely to increase power to detect differential isoform usage even in model organisms (Supplementary Fig. 7). Although the concept of superTranscripts is simple, it has the power to transform how studies of non-model organisms are performed as a multitude of the standard analytical tools and techniques can now be applied across all species.

## Materials and methods

### Datasets

To demonstrate how superTranscripts can be applied for visualization and differential transcript usage we used the public RNA-Seq dataset of human primary lung fibroblasts with an siRNA knock-down of *HOXA1* from Trapnell et al.[19] (GEO accession GSE37704). To demonstrate variant calling we sequenced RNA from genome in a bottle (GM12878) to a depth of 80 million 150bp paired-end reads using an Illumina NextSeq. The combined superTrancriptome for chicken was constructed using reads of chicken embryonic gonads from Ayers *et* al.[28] (SRA accession SRA055442).

### De novo transcriptome assembly and clustering

Trinity[20] version r2013-02-25 was used to assemble the Trapnell dataset into contigs using default options and a minimum contig length of two hundred. Contigs were then clustered by Corset[18] with the test for differential expression turned off (*-D 99999999*). The corset clustering and *de novo* assembled contigs were used as inputs to Lace to construct a superTranscript for each cluster. The superTranscript for each cluster was then assigned to a gene by aligning to the human reference superTranscriptome using BLAT with option -minIdentity=98.

### Read alignment

Reads were aligned to the genome or superTranscriptomes using the two-pass mode of STAR[21]. We used the STAR option ‐‐outSJfilterOverhangMin 12 12 12 to filter the output splice junctions. Junctions supported by 5 or more reads were used to define dynamic block positions. The annotation was created using Mobius, a python script in the Lace suite. Read alignments were visualized using Gviz[32]. Our R scripts for Gviz are provided at https://github.com/Oshlack/superTranscript_paper_code.

### Variant Calling

In order to detect variants using superTranscripts we ran the GATK Best Practice workflow for RNA-seq (https://software.broadinstitute.org/gatk/best-practices/rnaseq.php). We then filtered for heterozygous SNPs with at least 10 reads coverage. We used BLAT (options: -minScore=200 -minIdentity=98) to align all superTranscripts against the hg38 human genome. For superTranscripts that matched multiple genomic loci, we took the match with the highest score after excluding alternative chromosomes (denoted with “_alt” in the hg38 reference). We then constructed a chain file for liftOver (downloaded from http://genome.ucsc.edu/) and converted all superTranscript SNP positions into genome coordinates (scripts are available at https://github.com/Oshlack/superTranscript_paper_code). Finally, we excluded SNPs outside the Genome in a Bottle high confidence call regions and compared to the true positives reported by the Genome in a Bottle Consortium [22]. We also performed a genome-based approach to variant calling for comparison. For the genome-based analysis, the GATK workflow was followed using hg38 and subsequent steps were identical to the superTranscript analysis apart from translating coordinates to hg38 which was unnecessary. KisSplice version 2.4.0 was run with options, ‐‐max-memory 100GB -s 1 -k 41 ‐‐experimental. The genome positions for KisSplice SNPs were found by aligning the output file ending with “type_0a.fa”, against hg38 using STAR (options: ‐‐sjdbOverhang 73 ‐‐ sjdbGTFfile gencode24.gtf) and then running KisSplice2refgenome.

### Counting reads per bin

The featureCounts[25] function from Rsubread R package (v 1.5.0) was used to summarise the number of reads falling in a given genomic region. Reads were counted in paired end mode (-p) requiring that both ends map (-B). We allowed for reads overlapping multiple features (-O) assigning a fractional number of counts, 1/n, depending on how many features, n, the read overlapped (‐‐fraction). The same summarisation procedure was used in all cases.

### Differential isoform usage comparison

DEXseq[26] R package (v.1.5.0) was used to test for differential isoform usage. DEXSeq takes a table of block level counts from featureCounts and, by default, removes blocks with fewer than 10 counts summed across all samples. DEXseq then produces a per gene q-value as the probability that for a given gene there is at least one exon used differentially between conditions, controlling for multiple testing. SuperTranscripts were ranked on their q-values. For the de novo assembly analysis the use of superTranscripts was contrasted with Kallisto[10] and Salmon[11]. Kallisto and Salmon were run on the de novo assembled contigs using the default settings. Estimated counts per contig were then grouped into clusters using the same Corset clustering as was used by the superTranscripts. The count table was processed by DEXseq in the same way as the block counts table for superTranscripts. The “truth” was defined by mapping the reads to the human genome and quantifying differential isoform usage with DEXSeq based on the reference annotation. All genes that had a q-value < 0.05 were considered true positives whilst all genes with a q-value >0.9 were considered true negatives. Where multiple clusters were mapped to the same gene, the cluster with the lowest p-value was chosen and the others discarded. Clusters which mapped to multiple genes were removed from the analysis, and those genes found in a multi-gene cluster were removed from the list of true and false positives.

### Constructing a comprehensive chicken superTranscriptome

Ensembl and RefSeq references were downloaded for the chicken genome version galGal4 from UCSC on 24^th^ August 2016. Cufflinks transcripts were assembled using the gonad reads from Ayers et al.[28], mapped to galGal4 using TopHat[33] version 2.0.6. The reference and cufflinks assembled transcripts were then merged into loci based on genomic positions using the cuffmerge command. The resulting annotation was flattened and exonic sequence concatenated to create a genome-based superTranscriptome, similar to that described below for the human. To supplement these superTranscripts with de novo assembly, we first assembled all reads using Trinity. Trinty contigs were aligned against the genome-based chicken superTranscriptome using blat with options -minScore=200 -minIdentity=98. Contigs that aligned to a single genome-based superTranscript were clustered with it. Contigs matching two or more genome-based superTranscripts were discarded (to remove false chimeric transcripts[34]). Remaining contigs were clustered into genes based on their homology to human superTranscripts (using BLAT with options -t=dnax -q=dnax -minScore=200). Contigs that did not align to a gene, or those that aligned to multiple genes were removed. Lace was then run on the sequence in each cluster, containing genome-based superTranscripts and Trinity contigs.

In analysing the chicken superTranscriptome, we assessed the coverage from each constituent transcriptome, Ensmebl, RefSeq, Cufflink and Trinity, by aligning their sequence against the superTranscripts using BLAT with options -minScore=200 -minIdentity=98. We determined regions which were not present in the genomes galGal4 and galGal5 by aligning the superTranscripts against the chicken reference genome using BLAT with options -minScore=100 -minIdentity=98. Finally, we looked for regions with homology to human coding sequence by aligning the superTranscriptome against the Ensembl GRCh38 human protein sequence using BLAST[35] with options -evalue 0.00001 ‐ num_alignments 20. For a superTranscript region to be identified as novel protein coding sequence, we required it to be absent from the chicken genome, match a human protein sequence with BLAST e-value<10^−5^, only be annotated by a Trinity transcript and be 30bp or longer. Scripts used in the chicken superTranscript analysis are provided at https://github.com/Oshlack/superTranscript_paper_code.

### SuperTranscript construction for model organisms

When a reference genome was available we constructed superTranscripts by concatenating exonic sequence rather than using Lace. Doing so is more accurate as it does not rely on BLAT alignment or resolving loops in a splicing graph. The genome and annotation we used for human were taken from https://github.com/markrobinsonuzh/diff_splice_paper. As in Soneson et al.[27], the genome annotation was flattened, such that transcripts were merged into disjoint blocks. The sequence of each block was then extracted and concatenated for each flattened gene using the gffread -w command from the cufflinks suite. To annotate, we projected the genomic coordinates of transcripts onto the superTranscripts, then flattened the transcripts into blocks. The resulting human superTranscriptome, its annotation and the scripts used to create them are provided at https://github.com/Oshlack/superTranscript_paper_code.

## Acknowledgements

We would like to thank Paul Ekert and Stefanie Eggers for preparing and providing RNA-Seq from the genome in a bottle cell line. In addition we want to thank Ian Majewski, Jovana Maksimovic and Harriet Dashnow for feedback on the manuscript and Michael McLellan for his preliminary contribution. AO is funded by an NHMRC Career Development Fellowship APP1051481.

## Author Contributions

N.M.D and A.O. conceived the idea of superTranscripts and Lace. A.D.K.H. developed Lace and performed the differential isoform usage analysis. N.M.D. performed the SNP and chicken superTranscriptome analysis. All authors contributed to the writing of the manuscript.

